# Neuron-specific chromatin disruption at CpG islands and aging-related regions in Kabuki syndrome mice

**DOI:** 10.1101/2023.08.01.551456

**Authors:** Leandros Boukas, Teresa Romeo Luperchio, Afrooz Razi, Kasper D. Hansen, Hans T. Bjornsson

## Abstract

Many Mendelian developmental disorders caused by coding variants in epigenetic regulators have now been discovered. Epigenetic regulators are broadly expressed, and each of these disorders typically exhibits phenotypic manifestations from many different organ systems. An open question is whether the chromatin disruption – the root of the pathogenesis – is similar in the different disease-relevant cell types. This is possible in principle, since all these cell-types are subject to effects from the same causative gene, that has the same kind of function (e.g. methylates histones) and is disrupted by the same germline variant. We focus on mouse models for Kabuki syndrome types 1 and 2, and find that the chromatin accessibility abnormalities in neurons are mostly distinct from those in B or T cells. This is not because the neuronal abnormalities occur at regulatory elements that are only active in neurons. Neurons, but not B or T cells, show preferential chromatin disruption at CpG islands and at regulatory elements linked to aging. A sensitive analysis reveals that the regions disrupted in B/T cells do exhibit chromatin accessibility changes in neurons, but these are very subtle and of uncertain functional significance. Finally, we are able to identify a small set of regulatory elements disrupted in all three cell types. Our findings reveal the cellular-context-specific effect of variants in epigenetic regulators, and suggest that blood-derived “episignatures” may not be well-suited for understanding the mechanistic basis of neurodevelopment in Mendelian disorders of the epigenetic machinery.

## Introduction

The deposition and maintenance of epigenetic marks is one of the fundamental processes underlying development and homeostasis [1–4]. Reflecting this, the last decade has seen the emergence and rapid expansion of a group of developmental Mendelian disorders caused by disruptive *de novo* coding variants in genes encoding epigenetic regulators [5–8]. A characteristic of these disorders is the large number of affected organ systems, which exceeds what is seen in other Mendelian disorders [5]. This is consistent with the very broad expression of human epigenetic regulators across different tissues [9], and raises the question of whether the abnormalities in the different organs are driven by the same underlying chromatin disruption in the corresponding cell types. In other words: are the chromatin effects of variants in epigenetic regulators shared between different cell types, or are they primarily cell-type-specific?

There are two major reasons for why this question is important. First, from a basic biology standpoint, the answer would provide insights into whether the same variant, in the same epigenetic regulator, causes the same (or a highly similar) chromatin disruption in different cellular contexts. For example, is it the same genomic locations that exhibit changes in chromatin accessibility? The degree of such similarity would serve as an indicator of how much the intra- and extra-cellular environment dictate the specific effect that epigenetic regulators have on chromatin, and in what way. Second, from a practical standpoint, answering this question is important for understanding the potential of DNA-methylation-based “episignatures” [10, 11] to shed light into the pathogenesis of the disorders. While these episignatures are almost exclusively derived from whole blood, there is increasing emphasis on their capacity to pinpoint locations of chromatin disruption responsible for the dysfunction of other tissues/cell types [12]. In particular, because intellectual disability is the most prevalent phenotypic characteristic of these disorders, a major focus is on whether the same locations that are disrupted in blood also contribute to the abnormal neurodevelopment [12–14].

One of the prototypical Mendelian disorders of the epigenetic machinery is Kabuki syndrome (KS). Approximately 70% of cases of KS are caused by variants in either the histone methyltransferase *KMT2D* (KS type 1; KS1), or the histone demethylase *KDM6A* (KS type 2; KS2) [15, 16]. Several studies have recently investigated the molecular basis of KS, uncovering defects in different cell types [17–28]. In addition, KS was one of the first disorders for which episignatures were derived and shown to have diagnostic utility [29–31]. Therefore, KS is ideally suited for understanding the extent to which chromatin abnormalities caused by the disruption of the same epigenetic regulator are similar in different cell types.

Here, we investigate this using mouse models of KS1 and KS2, and profiling genome-wide chromatin accessibility in sorted neurons as well as sorted B and T cells from peripheral blood.

## Results

### Distinct disruption of chromatin accessibility in hippocampal neurons versus B and T cells in KS1 and KS2 mouse models

We performed ATAC-seq in order to characterize the genome-wide chromatin accessibility landscape in KS mice (both type 1 and type 2) and wild-type littermates (Methods). We examined sorted hippocampal neurons from the dentate gyrus (henceforth neurons) and sorted peripheral T cells (Methods). We also used previously generated ATAC-seq data from sorted peripheral B cells [18]. These cell types were chosen because: a) there is convincing evidence supporting their involvement in KS pathogenesis [18, 19, 22, 24, 26]; b) two of them (B and T cells) have similar functionality and common developmental origins, whereas neurons are functionally, morphologically, and developmentally distinct; c) they allow us to assess if blood episignature locations provide information about the epigenetic basis of abnormal neurodevelopment in KS.

We started with a differential accessibility analysis in neurons, after validating our detected ATAC peaks (Methods). Comparing 8 KS1 mice to 5 wild-type littermates (Methods), we discovered extensive chromatin disruption. Specifically, 20,683 (13.4%) of the 154,560 tested regulatory elements (ATAC peaks) exhibit altered accessibility (Supplemental Table 1; Methods). Focusing on promoter elements (+/− 2kb from the transcriptional start site; 19,013 peaks total), we estimated that 63.6% show disruption. We then found similar results in KS2, after comparing 4 mutant mice to 6 wild-type littermates (different from wild-type mice compared against the KS1 mutants). Specifically, we detected disruption of normal accessibility in 49,625 (31.7%) of the 156,309 tested neuronal regulatory elements (Supplemental Table 2; Methods), with 68.2% of promoter peaks affected.

We next asked if the same regulatory elements are disrupted in neurons and immune cells. We employed a previously developed method that uses conditional p-value distributions to evaluate the overlap between the results of genome-wide differential analyses, and has been shown to have increased power relative to more “naive” approaches ([18]; Methods). Specifically, here we started with one cell type (e.g. neurons or T cells) and asked if peaks disrupted in another cell type (e.g. B cells) have p-values that are overrepresented close to 0 in the starting cell type. If such an overrepresentation is present and exceeds what is expected by chance, it means that the same regulatory elements are preferentially disrupted in both cell types. In both KS1 and KS2, we discovered only a minor overlap between neurons and B cells genome-wide, whereas there is no statistically significant overlap when restricting to promoters (Figure 1a, b; Supplemental Figure S1a-d). In stark contrast, when comparing the disrupted elements in B cells to these in T cells, there is pronounced overlap both genome-wide and at the level of promoters; this is evident in both KS1 (Figure 1a, b; Supplemental Figure S1a, c) and in KS2 (Figure 1a, b; Supplemental Figure S1b, d).

**Figure 1.**
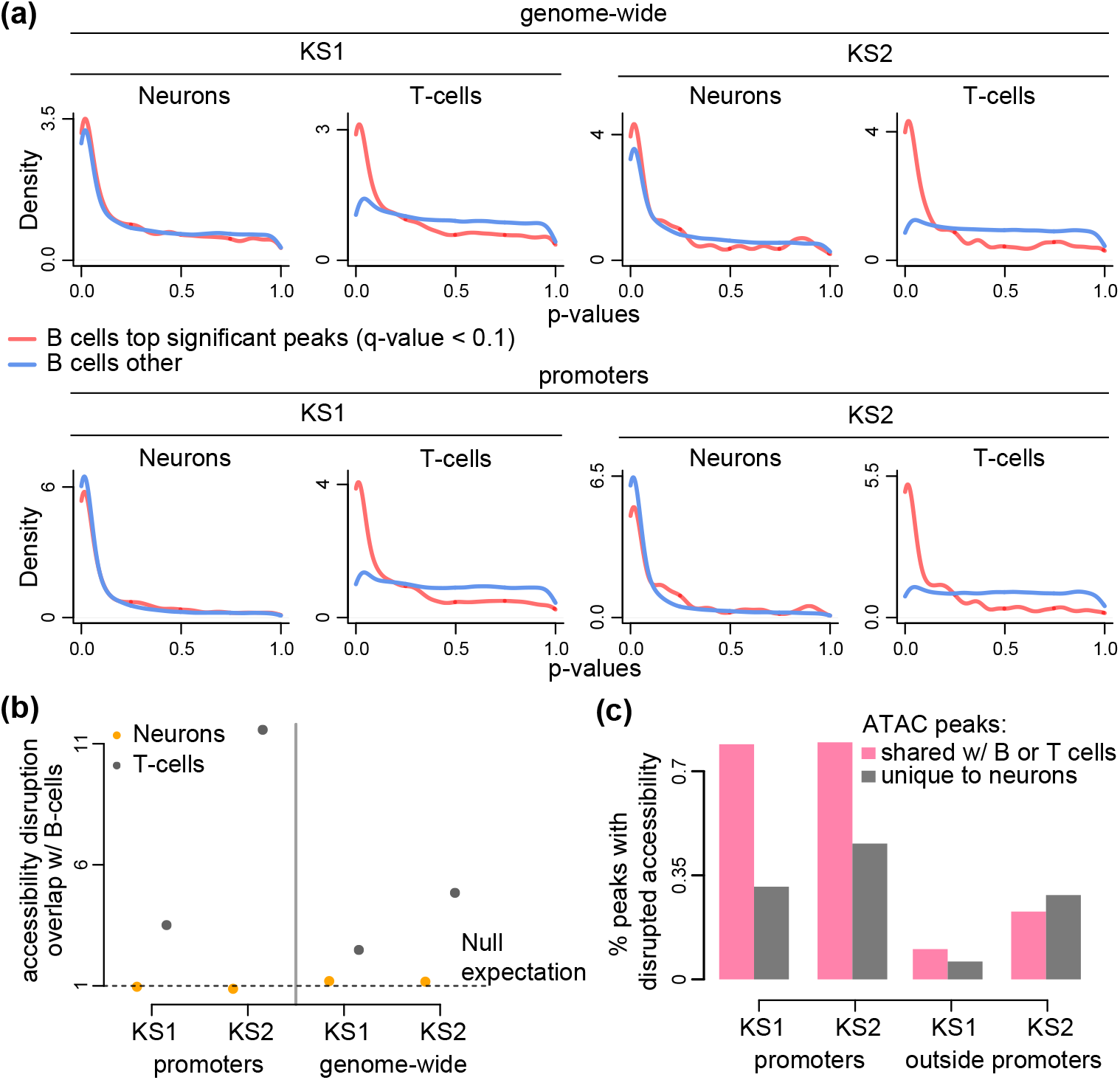
Evaluating whether neurons, B cells and T cells exhibit the same changes in chromatin accessibility in Kabuki syndrome types 1 and 2. **(a)** P-value distributions from mutant (KS1 or KS2) vs wild-type differential accessibility analyses in either neurons or T cells. Shown both for ATAC peaks genome-wide and for promoters only. The red densities correspond to ATAC peaks overlapping peaks disrupted in B cells (q-value *<* 0.1), whereas the blue densities correspond to ATAC peaks overlapping peaks not disrupted (but present) in B cells. **(b)** The ratio (y axis) of the proportion of ATAC peaks that overlap differentially accessible peaks in B cells and are also differentially accessible in either neurons (orange dots) or T cells (black dots), over the average proportion of ATAC peaks that overlap randomly sampled B cell peaks and are also differentially accessible in either neurons or T cells. The dashed horizontal line at 1 corresponds to the null expectation when there is no significant overlap between the ATAC peaks with disrupted chromatin accessibility in the two cell types tested. **(c)** The percentage of differentially accessible ATAC peaks in neurons (1 − *π*_0_; Methods), among ATAC peaks that are specific to neurons (grey bars) or ATAC peaks that overlap peaks that are also present in B or T cells (pink bars). Shown separately for promoter and non-promoter peaks in KS1 and KS2.

We note that all pairwise comparisons were performed after restricting to ATAC peaks detected in both cell types (Methods). This ensures that the greater overlap between B and T cells than between B cells and neurons is not driven by a greater overall similarity of the regulatory landscape. Related to this, we found that, with the exception of non-promoter elements in KS2, the neuronal chromatin disruption occurs predominantly at regulatory elements shared with either B or T cells, and not at elements uniquely active in neurons (Figure 1c; Supplemental Figure S2).

We next looked at mouse orthologs of human genes whose promoters are differentially methylated in KS1 patients versus controls in whole blood [29]; these are the promoters used to derive the KS1 episignature. Consistent with our other results, we found that these promoters are no more likely to be disrupted in neurons than randomly selected promoters (Supplemental Figure S3; Methods). By contrast, in B and T cells, these mouse ortholog promoters show clear evidence of preferential disruption compared to other promoters (Supplemental Figure S3), which suggests that the null result in neurons is not due to human-mouse differences.

### Chromatin accessibility alterations in Kabuki syndrome neurons, but not B or T cells, are concentrated at CpG island promoters

We set out to investigate the nature of the discrepancy between neurons and B/T cells. We focused on promoters, since their unambiguous association with the downstream gene can help connect that discrepancy to the underlying biology. A major distinction of mammalian promoters is between those that are associated with CpG islands, and those that are not [32]. We thus first asked if any systematic differences exist at this scale.

Ranking promoter ATAC peaks based on how strong the evidence for disruption of their accessibility is in KS1 neurons, we discovered a clear pattern: the stronger the evidence for accessibility disruption, the higher the probability of the promoter peak overlapping a CpG island (Figure 2a). Of the most confidently disrupted promoter peaks (smallest 10% of p-values), 94% overlap CpG islands, compared to 17% of the least confidently disrupted ones (highest 10% of p-values). In KS1 B or T cells, this relationship with CpG islands is completely absent (Figure 2a), despite the fact that the number of CpG islands containing ATAC peaks is – as expected – very similar in the three cell types (12,171 in B cells and 12,397 in T cells vs 11,377 in neurons). This indicates a fundamentally different localization of the most statistically robust chromatin alterations and sheds further light on our observation that broadly active regulatory elements show neuron-specific disruption. An almost identical picture is seen in KS2, with preferential disruption of CpG island promoters in neurons but not in B or T cells (Figure 2a). Further dissecting the neuronal chromatin disruption, we saw that, in KS1 as well as KS2 neurons, both of the main CpG island promoter classes [33] are disrupted: CpG island promoters bound by Polycomb Repressive Complex 2 (PRC2) and associated with tissue-specific genes (Methods), and – even more so – non-PRC2-bound CpG island promoters, that are associated with genes widely expressed across tissues (Figure 2b). We also found that, even though KMT2D and KDM6A are believed to promote transcriptional activity, disrupted CpG island promoters in mutant neurons almost always have increased accessibility relative to wild-type; Figure 2c).

**Figure 2.**
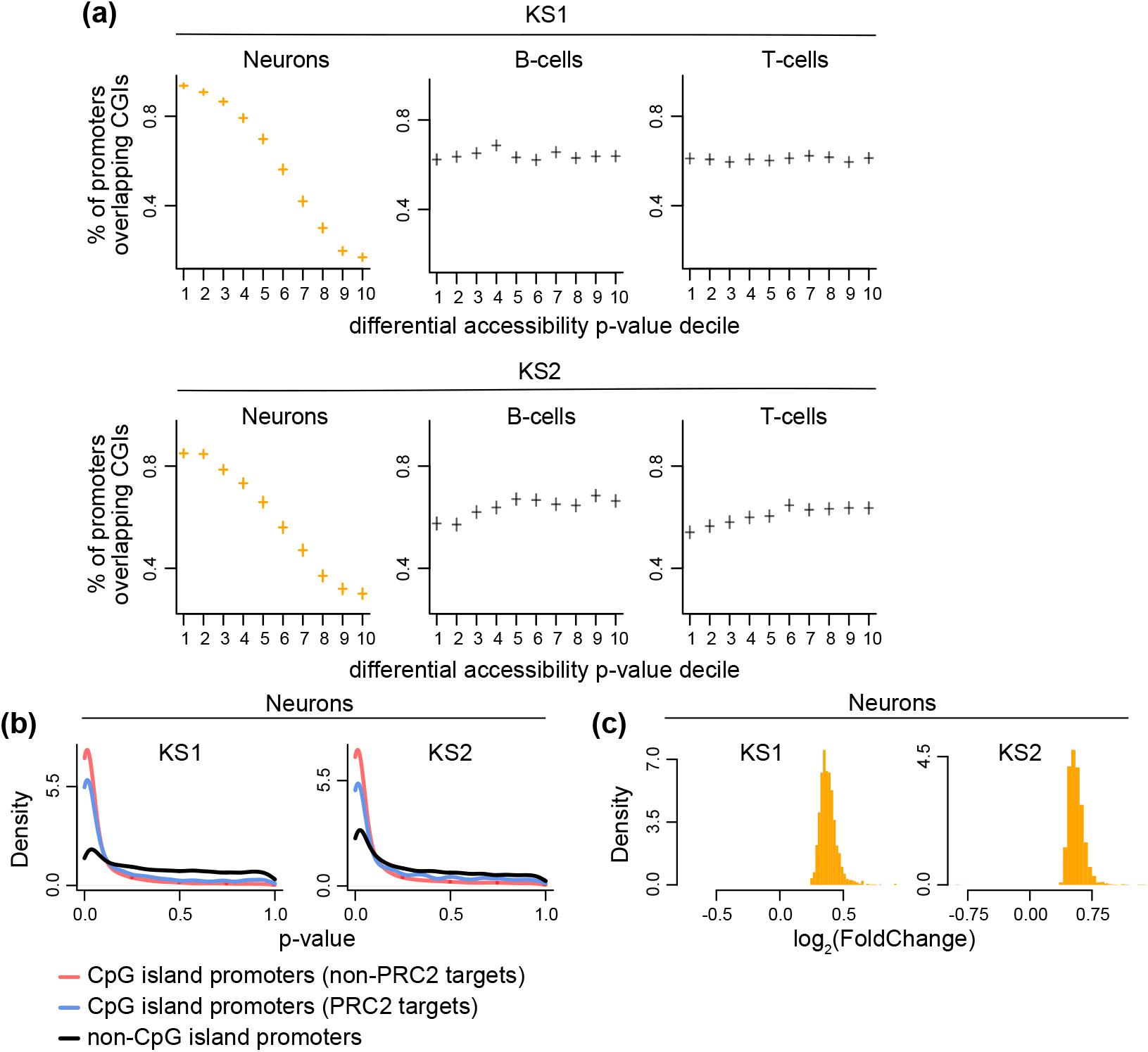
Assessing the relationship between promoter chromatin accessibility changes and promoter CpG-richness in neurons, B cells and T cells in Kabuki syndromes 1 and 2. **(a)** The proportion of promoter ATAC peaks overlapping CpG islands, stratified according to p-value decile. The horizontal lines correspond to these proportions across the deciles, and the vertical lines depict the associated 95% binomial confidence intervals. Shown separately for neurons, B and T cells, in KS1 (first row) and KS2 (second row). The p-values for these promoter peaks are derived from the corresponding differential accessibility analyses (Methods). **(b)** The p-value distributions of promoter peaks, stratified according to whether these peaks overlap CpG islands targeted by Polycomb Repressive Complex 2 (blue densities), overlap other CpG islands (red densities), or are located outside CpG islands (black densities). Shown for KS1 and KS2 neurons. **(c)** The distribution of the magnitude of accessibility changes (log_2_ (fold change)) of the most disrupted CpG-island-overlapping promoter peaks (bottom 10% p-value) in KS1 and KS2 neurons.

### Promoters of epigenetic machinery genes are preferentially disrupted in neurons but not B or T cells

It is known that the chromatin accessibility of CpG islands is related to the absence of DNA methylation and the presence of specific histone modifications [34–39]. We thus wondered if changes of such modifications are driving the observed changes in CpG island accessibility. While we did not directly measure histone modifications or DNA methylation, we reasoned that we can indirectly gain insight into this question by examining whether there is chromatin disruption at individual promoters of epigenetic machinery genes. If this is true in neurons, but not B or T cells, then that chromatin disruption could be driving the CpG island accessibility changes by causing changes in the expression of epigenetic machinery genes, and thus changes in histone modification and DNA methylation patterns at CpG islands. In both types of KS, we found that promoters of epigenetic machinery genes are preferentially disrupted compared to promoters of other genes in neurons (Figure 3a, b; Supplemental Tables 3, 4). However, this is not true in B or T cells (Figure 3a, b). The neuronal disruption is especially strong at a subset of epigenetic machinery genes whose human homologs have previously been shown to be highly intolerant to loss-of-function variation and coordinately expressed with one another in multiple tissues, and are thought to be particularly important in neurodevelopment (labeled as highly co-expressed EM genes in Figure 3a,b; Methods) [9]. In KS1, among the epigenetic machinery genes with disrupted promoters are *Kdm6a* (histone demethylase causing KS2; Figure 3c; *p* = 0.001), *Kmt2a* (histone methyltransferase causing Wiedemann-Steiner syndrome; Figure 3c; *p* = 0.004), *Tet3* (DNA demethylase causing Beck-Fahrner syndrome; *p* = 0.007), and *Chd1* (chromatin remodeler causing Pilarowski-Bjornsson syndrome; Figure 3c; *p* = 0.005). In KS2, epigenetic regulators with disrupted promoters range from *Kmt2d* (histone methyltransferase causing KS1; Figure 3d; *p* = 3.95 *·* 10*^−^*^6^) and *Suz12* (PRC2 subunit causing Imagawa-Matsumoto syndrome; Figure 3d; *p* = 4.08 *·* 10*^−^*^5^) to *Kat6a* (histone acetyltransferase causing KAT6A syndrome; Figure 3d; *p* = 0.001). These findings indicate broad epigenetic dysregulation in KS1 and KS2 that extends beyond what would be expected on the basis of the function of their respective causative genes, and could explain the disruption of CpG island chromatin. In addition, they may be relevant to the high degree of neurodevelopmental/neurological phenotypic overlap among the different Mendelian disorders of the Epigenetic Machinery [7].

**Figure 3.**
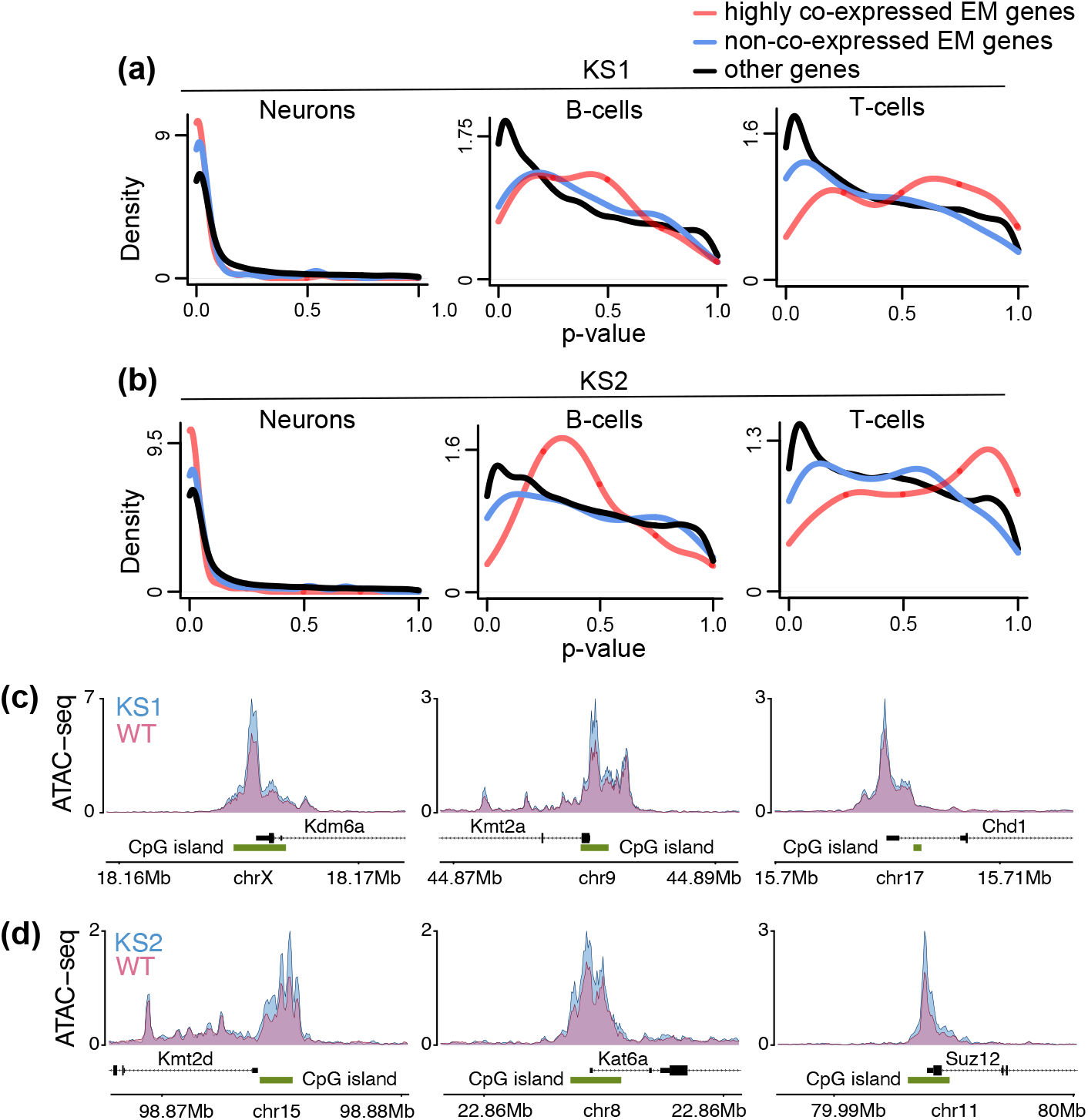
Investigating the chromatin disruption of promoters of epigenetic machinery genes in neurons, B cells and T cells in Kabuki syndrome types 1 and 2. **(a)** The p-value distributions of promoter peaks, stratified according to whether these peaks overlap promoters of highly co-expressed epigenetic machinery (EM) genes (red densities), non-co-expressed epigenetic machinery genes (blue densities), or promoters of other genes (black densities). See main text for details. Shown for neurons, B- and T-cells for KS1. **(b)** Same as **(a)** for KS2. **(c)** The normalized ATAC signal of 3 different epigenetic regulator genes in KS1 and wild-type neurons. **(d)** Like (c), but for KS2 neurons.

### Aging-related regions show preferentially disrupted chromatin accessibility in KS1 and KS2 neurons but not in B or T cells

Our findings yield insights into: a) the genomic distribution of the chromatin defects in KS1 and KS2; b) the cell-type selectivity of these defects. However, our findings thus far fall short of providing a link between the chromatin defects and the abnormal development known to be a component of KS. To obtain such a link, we first performed a pathway analysis of the genes with disrupted promoters in neurons (using Reactome; Methods). In both KS1 and KS2, the top pathways indicate disruption of various modes of epigenetic control (consistent with the results of the previous section) as well as disruption of other aspects of transcriptional regulation (Supplemental Table 5; Supplemental Table 6). However, these pathways are rather general. We thus searched for a characterization of the disrupted locations that better captures the abnormal developmental process in KS1 and KS2. To this end, we defined a set of regulatory elements whose chromatin state likely changes as development unfolds by restricting to ATAC peaks containing CpGs whose collective methylation status has been shown to be a highly accurate mouse age estimator – that is, an “epigenetic clock” (582 CpGs in total; Methods; Thompson et al. [40]). Apart from being biomarkers of biological aging, there is now emerging evidence that such epigenetic clocks track embryonic development as well [41, 42]. Using an approach that enables us to assess the collective disruption of a set of ATAC peaks even if not every individual peak is disrupted [43], we discovered that the chromatin accessibility of clock-CpG-containing regulatory elements is collectively disrupted in KS1 as well as KS2 neurons. This disruption is present not only at promoter age-associated elements (Figure 4a; lower values of the observed Wilcoxon rank-sum statistic than expected under the null distribution), but also at age-associated elements outside promoters (Figure 4a, d). By contrast, in B and T cells the chromatin accessibility of clock-CpG-containing regulatory elements elements is generally intact, both in KS1 and KS2 (Figure 4a).

**Figure 4.**
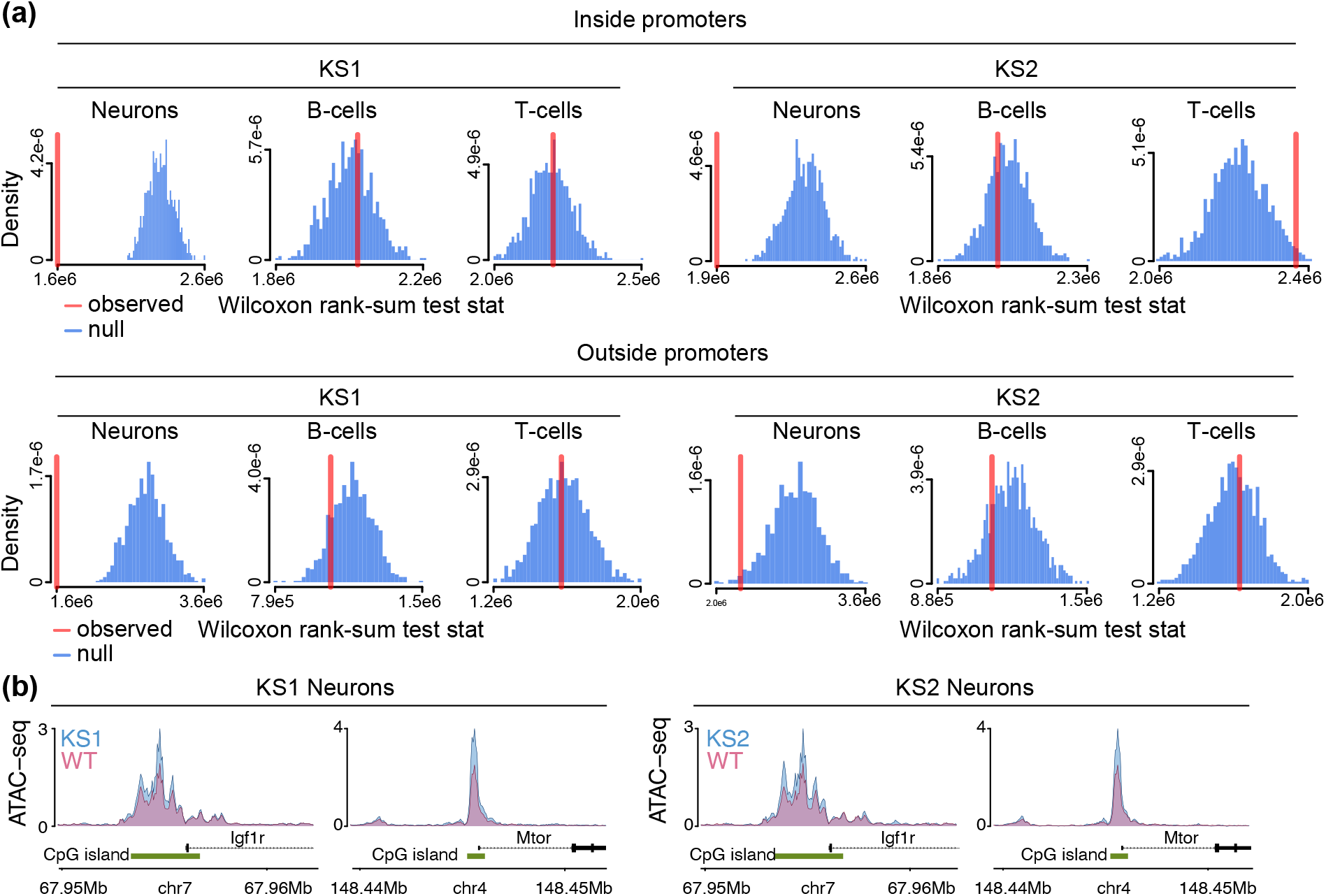
Investigating the chromatin disruption of aging-related regulatory elements in neurons, B cells and T cells in Kabuki syndrome types 1 and 2. **(a)** The observed value (red vertical line) of the Wilcoxon rank-sum test statistic obtained after comparing the differential accessibility p-values of aging-related ATAC peaks (that is, ATAC peaks which contain epigenetic clock CpGs) to the differential accessibility p-values of other ATAC peaks. Observed values lower than expected under the null (blue distributions) indicate the collective disruption of aging-related peaks. Shown separately for neurons, B cells, T cells, in KS1 and KS2, for ATAC peaks overlapping promoters and ATAC peaks outside promoters. The null distributions were derived by repeatedly (1000 times) sampling random sets of peaks, and calculating the Wilcoxon rank-sum test statistic after comparing the p-values of these randomly sampled peaks to the p-values of all other peaks. Each random set contained an equal number of peaks to the number of aging-related peaks. **(b)** The normalized ATAC signal of *Igf1r* and *Mtor*, two genes known to regulate lifespan-related pathways, in mutant and wild-type neurons. Shown for KS1 and KS2.

To further explore the link with regions involved in aging, we asked if genes implicated in lifespan-modulating pathways – such as signaling via the IGF1 receptor and mTOR signaling – have disrupted promoters. To do this in an unbiased manner, we examined genes in the mouse “longevity regulating pathway” as defined by KEGG (Kanehisa et al. [44]; Methods). None of the promoter ATAC peaks of these longevity genes contain any of the 582 clock CpGs, making this is an orthogonal assessment of the relationship between chromatin disruption in KS and aging-related regions. We found that in neurons, but not in B or T cells, promoters of longevity genes are collectively disrupted (Supplemental Figure S4). This is the case both in KS1 and KS2. Notable examples with disrupted promoters are *Igf1r* and *Mtor* (Figure 4b). Mutant neurons at the promoters of those two genes exhibit increased accessibility compared to wild-type, suggesting that the KS neurons are epigenetically “older” than their wild-type counterparts.

### The most significant chromatin alterations in B cells are in fact present, but very subtle, in neurons

All our investigations reveal substantial differences between the chromatin alterations in neurons and B/T cells. From a methodological standpoint, our results are based on p-values (although it is important to note that our approach has minimal reliance on arbitrary significance thresholds). As a final analysis, we returned to our overarching question of whether there is overlap between the chromatin disruption in immune cells and neurons, this time adopting an alternative approach. We asked whether the accessibility of the regions disrupted in B cells is altered in the same direction (increased or decreased) in neurons, more than expected by chance. In other words, we examined the concordance between the sign of the effect (log-fold change) in the two cell types, without taking the p-value into account [45–47]. This approach has greater sensitivity, and can thus answer whether regions disrupted in blood are also disrupted in neurons, but to a very small degree.

In all comparisons performed, we found statistically significant concordance in the direction of accessibility disruption (Figure 5a; Supplemental Figure S5). This includes the neuron vs B cell comparisons that previously showed no overlap based on p-value distributions. As expected, we did see that the concordance between B and T cells is always greater than the concordance between neurons and B cells (Figure 5a). Closer inspection of the behavior of neuronal peaks overlapping the most disrupted B cell peaks revealed that their accessibility alterations in neurons are of very small magnitude, which can explain why they do not stand out based on their p-values (pink curves; Figure 5b). However, the distribution of these magnitudes is different – centered away from 0 – from the distribution of the magnitude of the changes at elements not disrupted in B cells (pink vs blue curves; Figure 5b). This suggests that the neuronal disruption of the elements most disrupted in B cells is truly present, albeit very subtle.

**Figure 5.**
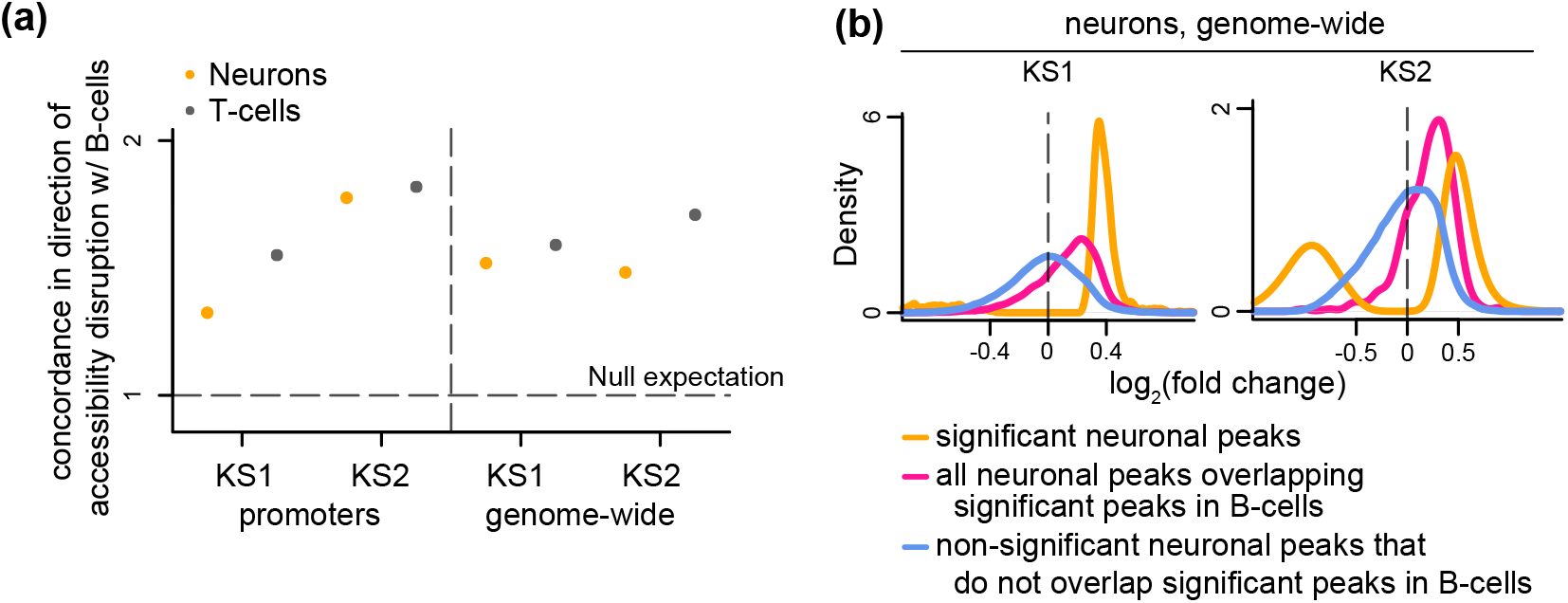
Evaluating whether chromatin accessibility changes occur towards the same direction in neurons, B cells and T cells in Kabuki syndrome types 1 and 2. **(a)** The ratio (y axis) of the proportion of ATAC peaks that overlap differentially accessible B cell peaks and exhibit accessibility changes in the same direction in neurons (orange dots) or T cells (black dots), over the average proportion of ATAC peaks that overlap randomly sampled B cell ATAC peaks and exhibit accessibility changes in the same direction in neurons or T cells (numerator and denominator in the ratio in Supplementary Figure S5). The dashed horizontal line corresponds to the null expectation when there is no significant concordance in the direction of the accessibility disruption (sign of log_2_(fold change)) between the two cell types tested. **(b)** The distribution of neuronal effect sizes (log2(fold changes)), shown for the significant disrupted neuronal ATAC peaks (orange), significantly disrupted neuronal peaks overlapping significantly disrupted B cell peaks (pink), and other neuronal peaks (blue). Significance is defined as q-value *<* 0.1.

### A small subset of regions exhibit changes in chromatin accessibility in all three cell types

Encouraged by the above, we suspected that it would be possible to define a set of regulatory elements with disrupted accessibility in all three cell types. Our ultimate goal was to derive, in a principled manner, a set of regions that can potentially comprise a blood-based biomarker that also accurately reflects the neuronal chromatin state. We employed a step-wise procedure. Starting from regions disrupted in B cells, we then selected the subset that are disrupted in neurons as well (10% FDR level). We then further narrowed these down by examining the accessibility of these regions in T cells, and selecting only those that are disrupted (10% FDR level). This procedure yielded 381 regions for KS1, and 91 regions for KS2 (Supplemental Tables 8, 9, 10, 12, 13, 14). The regions corresponding to the two KS types overlap much more than expected by chance: when restricting to promoter peaks and examining the genes immediately downstream (Supplemental Tables 7 and 11), there is an 66.3-fold enrichment of the KS1 genes in the set of KS2 genes (*p <* 2.2 *·* 10*^−^*^16^, Fisher’s exact test). This echoes previous results in blood and immune cells [10, 18]. Moreover, in each KS type, regions disrupted in both B cells and neurons are substantially more likely to be disrupted in T cells compared to regions only disrupted in B cells (Supplemental Figure S6). Taken together, these results reveal the existence of a small set of regions disrupted both in neurons and immune cells.

## Discussion

We have characterized, compared and contrasted the genome-wide chromatin alterations ensuing from genetic disruption of the histone methyltranferase *Kmt2d* and the histone demethylase *Kdm6a* in 3 different cell types: hippocampal neurons, and peripheral blood B and T cells. Our results paint a nuanced picture. On one hand, the most statistically significant chromatin alterations occur – mostly – at different genomic locations in neurons vs B/T cells (unsurprisingly, the alterations in the two immune cell types overlap substantially). On the other hand, the locations harboring the most significant chromatin disruption in B/T cells are in fact also disrupted in neurons, but the degree of this disruption is very small.

Broadly speaking, the differences between the cell types are compatible with three different scenarios. In the first scenario, when KMT2D and KDM6A exert their catalytic effects, the relative abundances of their interacting partners [48–51] are different in neurons versus immune cells. KMT2D and KDM6A are recruited to different locations as dictated by these abundances, and, as a result, when *Kmt2d* and *Kdm6a* are genetically disrupted the chromatin alterations also occur at different locations. In the second scenario, KMT2D and KDM6A exert their function at the same locations both in neurons and immune cells. However, other epigenetic regulators act after KMT2D and KDM6A to further modify the chromatin landscape in a different fashion in neurons versus immune cells. The chromatin alterations in this case may initially occur at the same locations, but are subsequently molded differently. In the third scenario, the chromatin alterations occur at a common progenitor cell type early during development. Then, subsequent chromatin changes occurring as part of neuronal and immune cell developmental programs [52, 53] ultimately give rise to what we observe in the mature cell types. The distinct chromatin alterations in this case are a reflection of distinct developmental paths. By contrast, the shared, subtle chromatin changes are remnants of the initial defects.

Notwithstanding the underlying explanation, we consider it surprising that a major difference between neurons and B/T cells is the extent to which the chromatin alterations are concentrated at promoter CpG islands of housekeeping genes; regulatory elements that have broad, non-cell-type-specific activity. Assuming that these chromatin alterations ultimately lead to abnormal levels of housekeeping gene protein products, this might also mean that the neuronal cellular phenotype is more severe, with fundamental processes affected. Also of potential mechanistic relevance is our finding that the chromatin disruption in neurons occurs preferentially at regulatory elements containing aging clock CpGs and at promoters of lifespan-associated genes. In other words, at locations that somehow track – and in some cases potentially regulate – the passage of biological time. This may represent a new link between the chromatin disruption in KS and the disruption of normal neural development/maturation. Our observation that, based on some of the accessibility changes, KS neurons may be epigenetically “older” than wild-type neurons fits with a previous report of precocious neuronal differentiation in KS1 [24] and warrants further functional exploration. The hyperactivation of the mTOR pathway that we infer has also recently been seen in cancer contexts upon *Kmt2d* as well as *Kdm6a* loss [54, 55]. We note, however, that a previous study reported a lower epigenetic age in patients with Kabuki syndrome, using the original Horvath epigenetic clock on blood samples [56]. The reasons for the discrepancy between these results is unclear but may include mouse-human differences, differences between cell types, as well as the fact that different variants have different effects on epigenetic age [56]. In addition, it should be pointed out that abnormal immune cell maturation is also known to be a component of the Kabuki syndrome phenotype [19, 22, 26]. Yet, the aging-associated locations are largely intact in B/T cells in our study, even though this clock was shown to function as an age predictor in multiple different mouse tissues (including blood; Thompson et al. [40]). In this respect, the chromatin basis of abnormal immune cell maturation appears distinct from the basis of abnormal neurodevelopment.

With respect to the very subtle neuronal chromatin alterations at the locations where the immune cells are most saliently disrupted, what our study does not answer is whether these subtle alterations are biologically significant in neurons. In other words, do alterations of such small magnitude have any effect on neuronal cellular function? Future work is needed to address this. It is also worth noting here that even the top significant neuronal alterations are of relatively small magnitude, with the majority of changes being much smaller than 2-fold. However, we anticipate that, collectively, these individually small changes can lead to the emergence of disease phenotypes [18, 57]. How small such chromatin changes can be and yet still have a biologically significant collective effect is, in our view, an important open question.

One limitation pertinent to all our results and interpretations is that the sorted cell populations we analyzed were defined by the expression of specific surface markers. However, it is now well-appreciated that cells expressing these markers represent a mixed population of cell sub-types [58–60]. Hence, it is possible that some of our conclusions would be different, had we analyzed such subtypes. For instance, we presently cannot exclude the existence of rare B or T cell sub-types whose chromatin disruption resembles that of neurons in terms of promoter CpG islands and aging-associated regulatory elements. Future single-cell analyses promise to provide an answer to this question, provided issues related to single-cell differential accessibility analyses are appropriately dealt with.

Finally, we emphasize that our goal when defining the sets of locations disrupted in all three cell types is not for these locations to replace those currently comprising existing KS episignatures; the latter have demonstrated their diagnostic utility [10, 11, 29]. However, if the objective is to somehow leverage the easily accessible epigenomic information from blood in order to draw conclusions pertinent to the chromatin basis of neurodevelopment in KS, we hypothesize that the locations we define in our current study may prove particularly useful. Future work is needed to test this, and evaluate if similar results hold for other MDEMs.

## Methods

### Mice

All experiments were performed in accordance with the National Institutes of Health Guide for the Care and Use of Laboratory Animals. All experiments received approval by the Animal Care and Use Committee of the Johns Hopkins University (protocol number: MO18M112). Both the KS1 (*Kmt2d*^+/*βGeo*^) and KS2 (*Kdm6a*^+/^*^−^*) mouse models and genotyping protocols have been previously described [18]. Mice for T cells are the same as reported in Luperchio et al. [18]. Neurons were obtained from 10-13 week old KS1, KS2 and wild-type mice.

### ATAC-seq: cell isolation and processing

#### B and T cells

B cells (CD19+) from peripheral blood were described in Luperchio et al. [18]. T cells were isolated at the same time as B cells. Specifically, flow through from binding and washing of CD19+ cells on Miltenyi separation columns was collected on ice. These CD19+ depleted samples were then spun and the supernatant was aspirated, resuspended and incubated with 2x (20uL) CD90.2+ microbeads (Miltenyi Biotec 130-121-278). CD90.2+ T cells were bound, washed and eluted per manufacturer protocol, then counted and aliquoted on ice for subsequent processing. Further processing of the T cell samples and ATAC sequencing was done in parallel with B cells as in Luperchio et al. [18]. A principal component analysis based on the ATAC signal profiles (after variance stabilization with the vst() DESeq2 function) shows essentially perfect separation between B and T cells (Supplemental Figure S7), as expected under efficient depletion of B (CD19+) cells prior to T cell isolation.

#### Neurons

Hippocampal neurons were isolated using a protocol adapted from the INTACT [61] and Omni-ATAC [62] protocols. Dentate gyrus was microdissected as previously described [24] and placed into ice cold 1xHBS (1x HBS-B, 1M sucrose, 500mM EDTA, 10% IGEPAL; 1x HBS-B was initially prepared as 6xHBS with 1MCaCl2, 1MMgAc, 1M Tris pH7.8, 1mM BME). Samples were homogenized in a 2mL glass dounce, pestle A until resistance went away (approximately 15 strokes) and pestle B until the tissue looked homogenized (approximately 8 strokes). Samples were then spun through a discontinuous iodixanol gradient. Gradients were made by diluting homogenized sample 1:1 with 50% iodixanol solution for a final concentration of 25% iodixanol (60% iodixanol solution + HBS), and overlayed on a 30%-35% iodixanol gradient in 15 mL conical tubes (30% iodixanol solution: 1xHBS, 30% iodixanol solution, 160mM sucrose; 35% iodixanol solution: 1xHBS, 35% iodixanol solution, 160mM sucrose). Gradients were spun for 5000g 20min 4’ with no break in swinging bucket centrifuge. Nuclei were collected at the 30-35% interface, transferred to a new tube and diluted with 1xHBS. Samples were spun for 10min 4’ 1000g, aspirated, then resuspended in 200uL 1xHBS. Nuclei were then stained for NeuN (NeuN-488, EMD Millipore MAB377X) and incubated 30min in dark. DNA was counterstained with Hoechest 33342 and sorted on NeuN+ to collect neurons (to avoid glia). 5000 nuclei were collected for most samples, though some variation was present. Additional processing, library preparation, and ATAC sequencing was done as for B/T cells following the protocol described in Luperchio et al. [18], adjusted for amplification cycles for ATAC.

### ATAC-seq: mapping and peak calling

Raw ATAC-seq data were processed as follows. Sequencing reads from the fastq files were mapped to the mouse genome (assembly mm10) using bowtie2 [63], duplicate reads were removed with Picard (via the “MarkDuplicates” function), and mitochondrial reads were removed with samtools [64]. Then, separately for each cell type, we identified genotype-specific peaks as follows. First, we merged the bam files corresponding to samples of each genotype (KS1, KS2, wild-type; one sample from each mouse) using the “merge” function from samtools. This created 3 “meta-samples”, each one corresponding to a single genotype (the wild-type mice from both the KS1 cohort and the KS2 cohort were merged into the same “meta-sample”). We subsequently called peaks from these 3 meta-samples with MACS2 [65], with the “keep-dup” parameter set to “all”.

To orthogonally validate the peak locations and ATAC signal intensities we detected in neurons, we first examined ATAC-seq data from the post-natal (day P0) mouse forebrain (since the hippocampus is part of the forebrain). These data were generated by Gorkin et al. [53] as part of the ENCODE project and can be obtained from https://hgdownload.soe.ucsc.edu/gbdb/mm10/encode3/atac/. They consist of a set of genomic regions, with each region having a corresponding ATAC signal score. We found that 98% of KS1, 98.5% of KS2, and 97.8% of the wild-type neuronal peaks overlap ENCODE regions with score greater than 0. This supports the validity of our peak locations. To then assess the validity of the ATAC signal intensity at these locations, we restricted to ENCODE regions with a score greater than 0, and stratified them according to whether they: a) overlap at least one of our ATAC peaks at promoters; b) overlap at least one of our ATAC peaks outside promoters; c) do not overlap any of our peaks. Examining the scores of the stratified regions, we found that, in each of the genotypes, promoter-overlapping regions have strong signal, and non-promoter overlapping regions have weaker (as expected), but still pronounced signal (Supplemental Figure S8a). By contrast, ENCODE regions that do not overlap any of our peaks show very weak signal (Supplemental Figure S8a).

As a final validation, we compared the ATAC peaks we detected in our samples to the peaks detected by Su et al. [66] (GSE82010). This study is well-suited as a comparison, as it also examines the dentate gyrus of the mouse hippocampus. To ensure a robust comparison, we downloaded the raw fastq files corresponding to the wild-type mice, and processed them in an identical manner as our own fastq files. We then called peaks after creating a single wild-type “meta-sample” by merging all bam files together. We found that, the vast majority of the ATAC peaks identified in Su et al. [66] are detected in our samples as well (96.4% overlap wild-type peaks, 97.4% overlap KS1 peaks, and 90.4% overlap KS2 peaks; Supplemental Figure S8d).

Together, these results provide strong evidence that our neuronal ATAC-seq peaks represent true neuronal regulatory elements, in all three of the genotypes.

### ATAC-seq: differential accessibility analysis

For neurons and T cells, the differential accessibility analyses were performed separately, as follows. First, we unionized the 3 sets of peaks called from the 3 meta-samples (where each metasample corresponded to one genotype; see previous section) and excluded peaks which had at least 1 base-pair overlap with ENCODE blacklisted regions (obtained from https://github.com/Boyle-Lab/Blacklist/archive/v2.0.zip; Amemiya et al. [67]). This yielded a common (across genotypes) set of peaks for each cell type: 172,558 peaks in neurons and 93,446 peaks in T cells [18]. Using this common set of peaks, for each differential analysis (e.g. KS1 vs wild-type in neurons), we created a peaks-by-samples matrix by counting the number of reads from each sample that map to each peak with the featureCounts() function from the Rsubread R package [68] (requiring an overlap of at least 3 base pairs between a read and the corresponding peak location). We specified the parameters of featureCounts() so as to exclude reads that map to multiple locations and reads belonging to chimeric fragments. The resulting counts served as the entries of the peaks-by-samples matrix. Using the peaks-by-samples matrix, we excluded peaks with median count (across samples) less than 10. Finally, we performed the differential analysis with DESeq2 [69], after adjusting for unknown confounding variables estimated via Surrogate Variable Analysis [70]. For B cells, we obtained the differential accessibility results from Luperchio et al. [18], but also included peaks that had been excluded from the analysis therein due to adjusted p-values set to NA after independent filtering by DESeq2.

To validate our differential accessibility results in KS1 neurons, we stratified our ATAC peaks based on whether they overlap regions where KMT2D has been previously shown to bind in ht22 cells (via ChIP-seq; Carosso et al. [24]). This is a mouse hippocampal neuronal cell line, and thus provides a reasonable comparison. We found that ATAC peaks overlapping KMT2D ChIP-seq peaks have a higher probability of exhibiting disrupted accessibility compared to other ATAC peaks, as evidenced by the corresponding p-value distributions (see also next section). This is true both for ATAC peaks at promoters and ATAC peaks outside promoters (Supplemental Figure S9). We also found strong overlap between the regions with disrupted neuronal accessibility in KS1 and KS2 (Supplemental Figure S10).

### Comparing the chromatin disruption between different cell types

Using the results of the differential accessibility analyses, we compared the chromatin disruption between cell types using the approach developed in Luperchio et al. [18] (where it was applied for comparisons between genotypes). First, we obtained the top differentially accessible ATAC peaks in B cells. Then, we obtained neuronal or T cell peaks that have an overlap of at least 1 base pair with the B cell differential peaks, and examined the distribution of the p-values corresponding to these peaks. These p-values were obtained from the differential accessibility analysis in neurons or T cells, respectively. From that p-value distribution, we estimated the proportion of these peaks that are differential in neurons or T cells (in addition to being differential in B cells) using Storey’s method [71, 72]. This proportion (1 *− π*_0_) corresponds to what we often refer to in the main text as the overlap between the two cell types. We computed *π*_0_ using the pi0est() function from the “qvalue” R package, with the “pi0.method” argument set to “bootstrap”. To test whether the resulting overlap is statistically significant, we derived a null distribution by repeatedly (10,000 times) sampling random sets of B cell peaks, and estimating the neuronal or T cell *π*_0_ from these peaks in an identical manner as for the B cell differential peaks. The number of peaks contained in each randomly sampled set was equal to the number of differential B cell peaks.

The above procedure requires us to choose initial thresholds in order to define the top differentially accessible peaks in B cells. For Figure 1, we used a q-value threshold of 10%. For KS2, the bootstrap method occasionally failed to yield an estimate for the null *π*_0_ during the permutations, likely due to the paucity of large p-values. Hence, we used the default method implemented in the pi0est() function but with the “lambda” parameter set to 0.5, because we have empirically observed that this value generally yields estimates consistent with these of the bootstrap method. We also sought to verify that our findings are not dependent on the specific thresholds used. We performed the same overlap analysis, but this time starting with the differential accessibility results in neurons or T cells, instead of B cells. That is, we defined differential peaks in neurons (q-value *<* 0.1), and T cells (q-value *<* 0.1). We subsequently examined the p-value distribution of these peaks in B cells, and estimated *π*_0_ as before. Our results unambiguously recapitulate our original findings: there is much greater overlap between the chromatin disruption in B and T cells, than between B cells and neurons (Supplemental Figure S1). In addition, there is no statistically significant overlap between B cells and neurons.

Finally, for our analysis of the concordance in the direction of accessibility disruption between cell types, the null distributions were derived as follows. We repeatedly (1,000 times) sampled random sets of B cell peaks. The sampling was done so that, within each random set: a) the balance of peaks with increased vs decreased accessibility in the mutant vs wild-type mice was the same as in the set of B cell differential peaks (see above for defining differential peaks); b) the total number of peaks sampled was equal to the number of B cell differential peaks. For each random set of B cell peaks, we obtained overlapping neuronal or T cell peaks and examined the proportion of these peaks whose accessibility changes in the same direction as in the B cell differential peaks. These random proportions formed the null distribution.

### Mouse promoter coordinates

We defined promoters as regions +/− 2kb from the transcriptional start site. We obtained coordinates of such promoter regions in mm10 using the promoters() function from the EnsDb.Mmusculus.v79 R package, with the “upstream” and “downstream” parameters set to 2000.

### Mouse orthologs of human episignature genes

We obtained the coordinates of differentially methylated positions (DMPs) in KS1 patients vs controls from Aref-Eshghi, Schenkel, et al. [29]. We then used the EnsDb.Hsapiens.v75 R package to obtain the coordinates of human promoters in hg19, and restricted to promoters that contain at least one DMP. To minimize false positive associations between promoters and DMPs, here we defined promoters as regions +/− 500bp from the transcriptional start site. We mapped the ENSEMBL id’s of the genes corresponding to these promoters to their mouse ortholog ENSEMBL id’s using the biomaRt R package (with the “mmusculus homolog orthology confidence” parameter set to 1). To perform the analysis, we then obtained gene-level p-values for these genes as described in the “Pathway analysis” section below.

### CpG islands

We obtained coordinates of CpG islands in the mouse genome (mm10 assembly) using the ucscTableQuery() function from the “rtracklayer” R package. We defined ATAC peaks that overlap CpG islands as peaks with at least 1 overlapping base pair.

As expected, peaks overlapping CpG islands have higher read counts compared to peaks that do not. Since the number of read counts can be inversely related to the uncertainty around log fold-change estimates, we sought to verify that the strong quantitative relationship between probability of CpG island overlap and differential accessibility p-value (Figure 2) is not driven by the higher read counts of CpG-island-overlapping peaks in the setting of lower overall counts in neurons (across-peak median average normalized read count = 151.4 and 124.2 in KS1 and KS2, respectively) versus B (across-peak median average normalized read count = 368 and 400.4 in KS1 and KS2, respectively) or T cells (across-peak median average normalized count = 296.2 and 302.8 in KS1 and KS2, respectively). We downsampled the bam files corresponding to the B and T cell samples to 25% of the initial number of reads, generated new peaks-by-samples matrices using the same procedure as before, and performed a new differential accessibility analysis. We found that, even with overall counts similar to these in neurons, there is still no relationship between probability of CpG island overlap and differential accessibility p-value in B or T cells; this is true both in KS1 (1*−* ratio of deviance to null deviance = 1.58 *·* 10*^−^*^5^ in B cells and 1.57 *·* 10*^−^*^5^ in T cells before downsampling vs 7 *·* 10*^−^*^5^ and 10*^−^*^3^ after downsampling) and KS2 (1*−* ratio of deviance to null deviance = 0.003 in B and T cells before downsampling vs 8 *·* 10*^−^*^4^ in B cells and 2 *·* 10*^−^*^4^ in T cells after downsampling).

To identify mouse CpG islands bound by the Polycomb Repressive Complex 2 (PRC2), we first identified locations bound by EZH2 – the catalytic subunit of the PRC2 complex – in the human genome. To do this, we used the ucscTableQuery() function from the “rtracklayer” R package to query the UCSC Table Browser. Specifically, from the “Txn Factor ChIP” track (which belongs to the “Regulation” group) we obtained the “wgEncodeRegTfbsClusteredV3” table. This provided us with a set of genomic locations and their coordinates, with each location having an associated score. The maximum possible score was 14. That score represents the number of EZH2 ChIP-seq peaks detected in that location, after uniform processing of ENCODE ChIP-seq experiments from 14 cell lines (H1-hESC; embryonic stem cells, HeLa-S3; cervical carcinoma, HMEC; mammary epithelial cells, HSMM; skeletal muscle myoblasts, NH-A; astrocytes, NHDF-Ad; dermal fibroblasts, NHEK; epidermal keratinocytes, NHLF; lung fibroblasts, Dnd41; T cell leukemia with Notch mutation, GM12878; lymphoblastoid, HepG2; hepatocellular carcinoma, HSMMtube; skeletal muscle myotubes differentiated from the HSMM cell line, HUVEC; umbilical vein endothelial cells, and K562; lymphoblasts). To minimize the inclusion of potential false positives in our analysis, we restricted to locations having at least 5 EZH2 peaks (2,776 regions). We subsequently identified human genes whose promoters overlap these locations (that is, have at least 1 base pair in common) using the promoters() function from the “EnsDb.Hsapiens.v75” R package. Then, we obtained mouse orthologs of these human genes using the “biomaRt” R package. We only selected high-confidence ortholog pairs, by setting the “mmusculus homolog orthology confidence” parameter equal to 1. This yielded a total of 1,515 mouse orthologs. Finally, we identified mouse CpG islands that overlap the promoters (+/−2 2kb from the TSS) of these mouse orthologs.

### Normalized ATAC signal plots

The BAM files corresponding to the genotype-specific “meta-samples” were indexed (using samtools) and converted to bigwig files using deeptools [73] (see https://deeptools.readthedocs.io/en/develop/). Reads were CPM normalized also using deeptools (bamCoverage –normalizeUsing CPM). Mouse Transcript annotations as well as CpG island annotations (both from mm10) were obtained from UCSC. All plots were then generated using the karyoploteR R package. We plotted regions between 5000bp upstream and 8000bp downstream of the selected promoters using the plotKaryotype() function. The normalized ATAC signal tracks were plotted with the kpPlotBigWig() function, from the bigwig files. Gene tracks were added by addGeneNames() and mergeTranscripts() and added to the plot by kpPlotGenes() function. CpG islands overlapping the promoter regions were added to the plot by using kpPlotRegions(). Labels and text positions were determined by “r0” and “r1” parameters in karyoploteR.

### Epigenetic machinery genes

Epigenetic machinery (EM) genes and their co-expression status (highly co-expressed vs non-co-expressed) were obtained from [9]. Therein, it was shown that highly co-expressed EM genes are these whose expression levels are correlated (across individuals) with those of many other EM genes in multiple different tissues. Non-co-expressed EM genes are these whose expression is not correlated with the expression of other EM genes.

### Pathway analysis

The differential accessibility analyses provide results at the peak level. To perform a pathway analysis, we obtained a single p-value for each gene by selecting the smallest p-value out of the p-values corresponding to the peaks that overlap the gene’s promoter (or the gene’s multiple promoters, in cases where there are more than one annotated TSS’s). We defined differential genes as those with p-values in the bottom 5%. We subsequently used these differential genes and the Reactome pathway annotations [74] to identify disrupted pathways with the “goseq” R package [75].

### Mouse epigenetic clock CpGs and mouse longevity genes

We obtained mouse epigenetic clock CpGs from Thompson et al. [40]. Specifically, we used the “elastic net clock”, which is comprised of 582 CpGs. This clock is a linear model whose predictors are the methylation values of these 582 CpGs. It was shown in Thompson et al. [40] that the age predicted by this clock correlates strongly with chronological age across a diverse set of tissues, including blood and cerebral cortex.

To examine the chromatin disruption at promoters of genes associated with lifespan regulation in mammals, we used the KEGGREST R package to obtain genes in the “Longevity regulating pathway” (pathway: mmu04213). We then analyzed peaks within promoters (+/− 2kb) of these genes. For Figure 4b, the gene-level p-values were obtained as described in the “Pathway analysis” section above.

### Identification of regulatory elements disrupted in all three cell types

In both KS1 and KS2, we adopted the following stepwise strategy. We started by obtaining peaks disrupted in B cells (q-value *<* 0.1). Among these genes, we then seelcted peaks disrupted in neurons as well (at 10% FDR level) using the qvalue() function from the “qvalue” R package [71]. Finally, from the resulting peaks, we selected these disrupted in T cells (10% FDR), again using the qvalue() function.

## Data and Code Availability

ATAC-sequencing data can be found under GEO accession GSE239339 (superseries containing the neuron and T cell data as subseries). Code for all analyses can be found in https://github.com/hansenlab/mdem_neuron_paper

## Acknowledgements

HTB and TRL were supported by a grant from the Louma G Foundation. HTB was also supported by the Icelandic Research Fund (#217988, #195835, #206806) and the Icelandic Technology Development Fund (#2010588). Research reported in this publication was supported by the National Institute of General Medical Sciences of the National Institutes of Health under award number R01GM121459 and CZF2019-002443 from the Chan Zuckerberg Initiative DAF, an advised fund of Silicon Valley Community Foundation. The funders had no role in study design, data collection and analysis, decision to publish, or preparation of the manuscript.

## SUPPLEMENTARY MATERIALS

### 1 Supplemental Figures

**Supplementary Figure S1.**
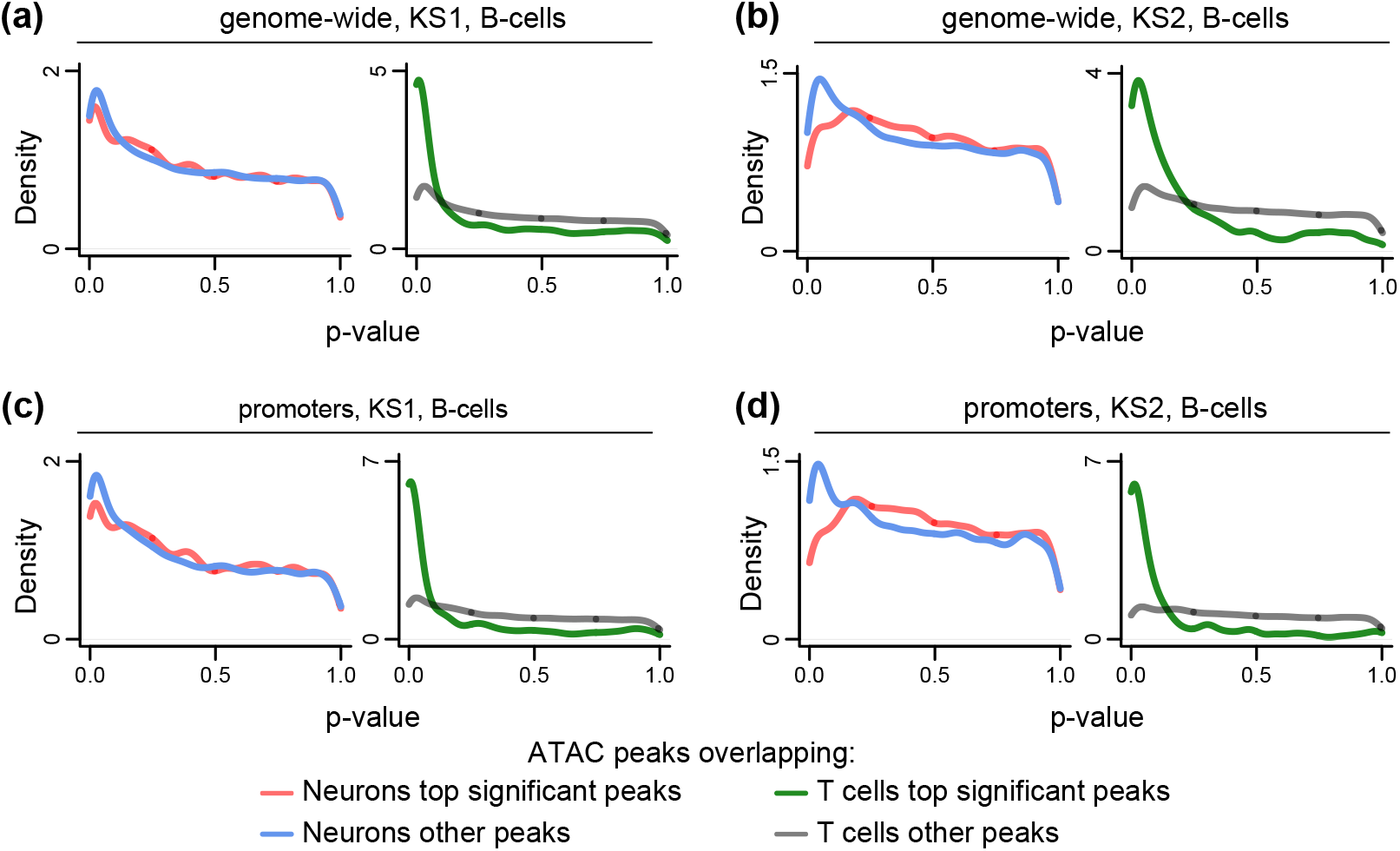
Evaluating whether neurons, B cells and T cells exhibit the same changes in chromatin accessibility in Kabuki syndrome types 1 and 2; alternate conditioning. **(a) -(d)** Like Figure 1a, but where the x-axis corresponds to p-values from the mutant (KS1 or KS2) vs wild-type differential analysis in B cells, and the conditioning (i.e. the stratification into red vs blue or green vs gray densities) is done either based on which peaks overlap peaks with disrupted accessibility in neurons (red vs blue), or based on which peaks overlap peaks with disrupted accessibility in T cells (green vs gray).

**Supplementary Figure S2.**
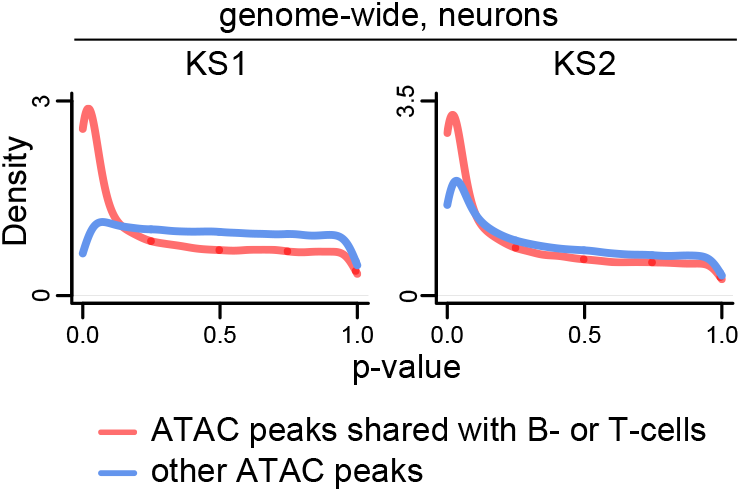
Evaluating the relationship between probability of differential accessibility in KS1 and KS2 neurons and neuron-specific vs broad peak activity. The distributions of p-values from the KS1 and KS2 mutant vs wild-type differential accessibility analyses in neurons, stratified according to whether the ATAC peaks are specific to neurons (blue densities) or overlap peaks that are also present in B or T cells (red densities).

**Supplementary Figure S3.**
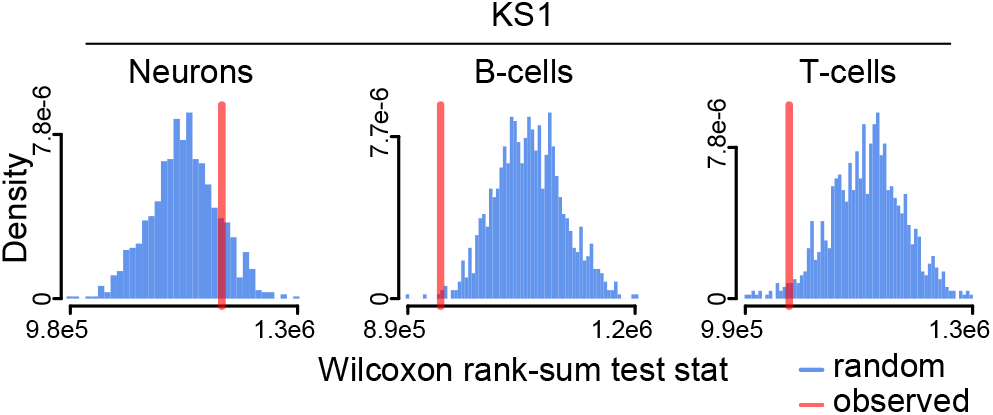
Investigating the chromatin disruption of promoters of mouse orthologs of human episignature genes. The observed value (red vertical line) of the Wilcoxon rank-sum test statistic obtained after comparing the p-values corresponding to mouse orthologs of human genes whose promoters contain CpGs differentially methylated between KS1 patients and controls in whole blood to p-values corresponding to all other genes. See Methods for details. Observed values lower than expected under the null (blue distributions) indicate the collective disruption of aging-related peaks. The observed value (red vertical line) was obtained by comparing The null distributions were obtained by repeated random sampling from all genes to obtain a null gene set of equal size to the set of episignature homolog genes and compare it to all other genes.

**Supplementary Figure S4.**
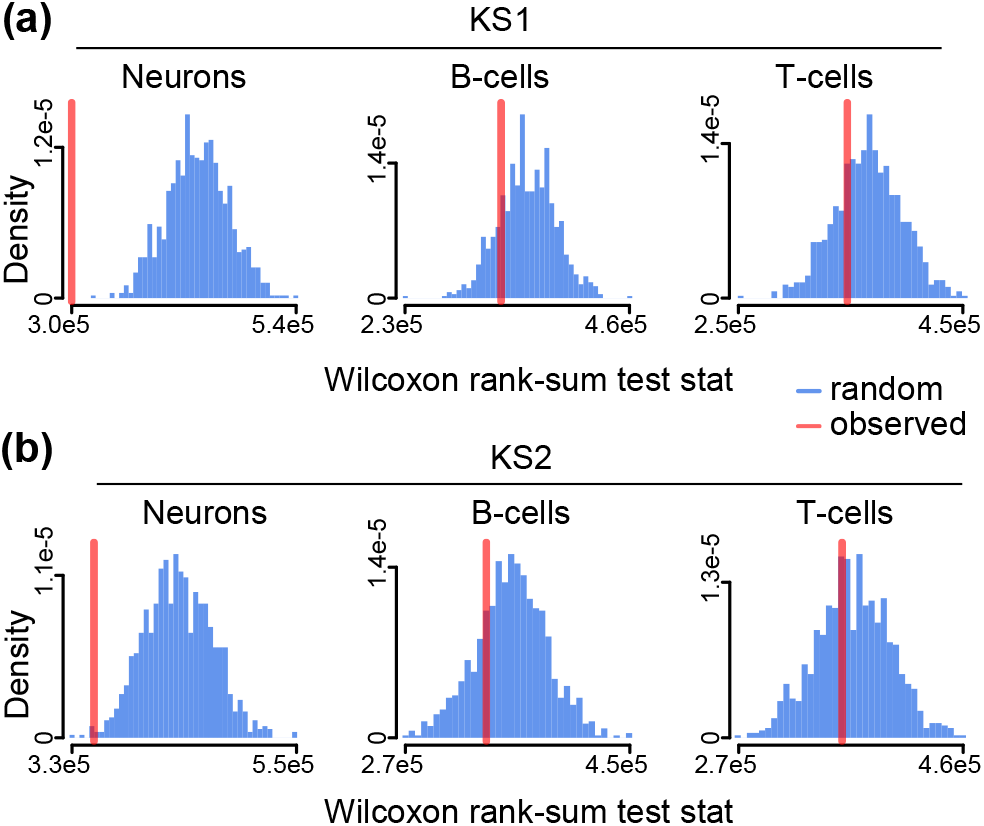
Investigating the chromatin disruption of promoters of longevity-associated genes in neurons, B cells and T cells in Kabuki syndrome types 1 and 2. **(a)-(b)** Like Figure 4a, but for ATAC peaks overlapping promoters of genes in the mouse KEGG “longevity regulating pathway”. Shown for KS1 and KS2. See Methods for details.

**Supplementary Figure S5.**
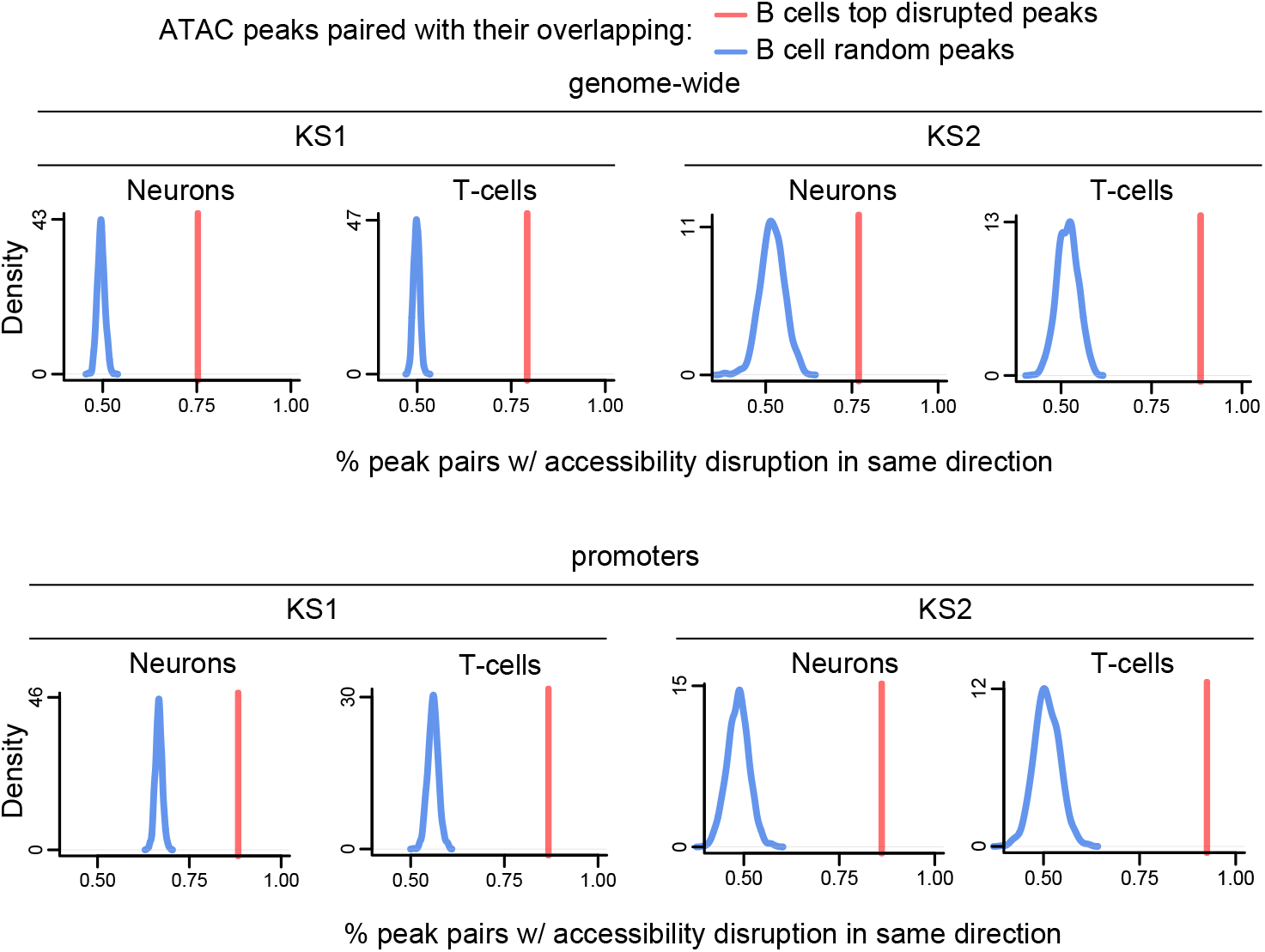
Evaluating whether chromatin accessibility changes are towards the same direction in neurons, B cells and T cells in Kabuki syndrome types 1 and 2; null distributions vs observed values. The red vertical lines depict the observed percentage of ATAC peaks (either in neurons or T cells) that overlap the top significantly disrupted peaks in B cells (qvalue *<* 0.1) and show accessibility changes towards the same direction. The blue densities correspond to null distributions, obtained by repeatedly (1000 times) sampling a random set of B cell peaks and computing the percentage of overlapping neuronal peaks that show accessibility changes towards the same direction. The sampling is performed so as to ensure that, among the random peaks, the balance of peaks with increased vs decreased accessibility in mutants vs wild-type is the same as among the top disrupted B cell peaks.

**Supplementary Figure S6.**
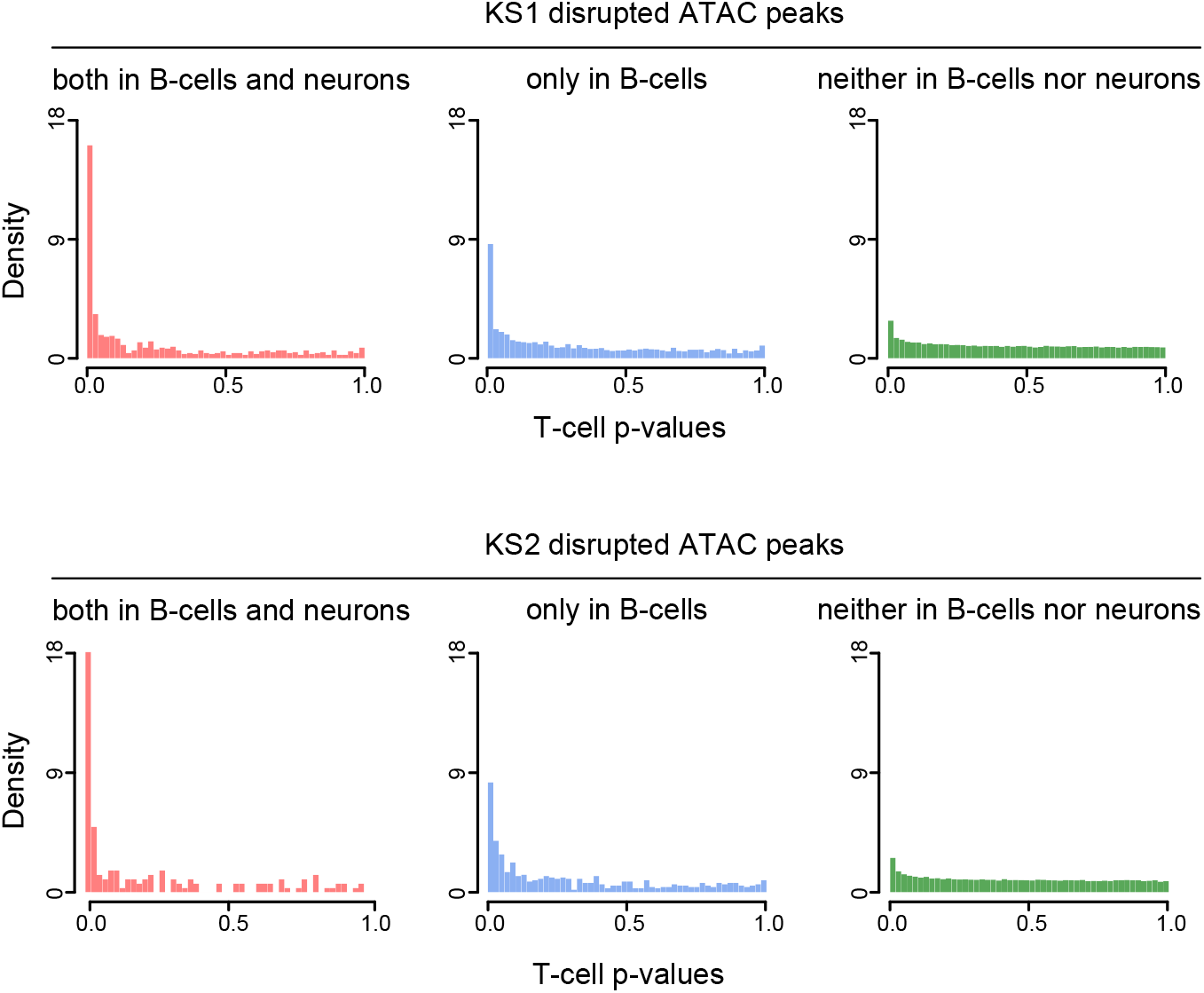
Evaluating whether neuronal chromatin disruption can distinguish peaks disrupted in T cells on top of information provided by disruption in B cells. The p-value distributions of T cell ATAC peaks that overlap peaks that are: differential in both B and neurons; differential in B cells but not in neurons; neither differential in B cells nor neurons. Depicted separately for KS1 and KS2.

**Supplementary Figure S7.**
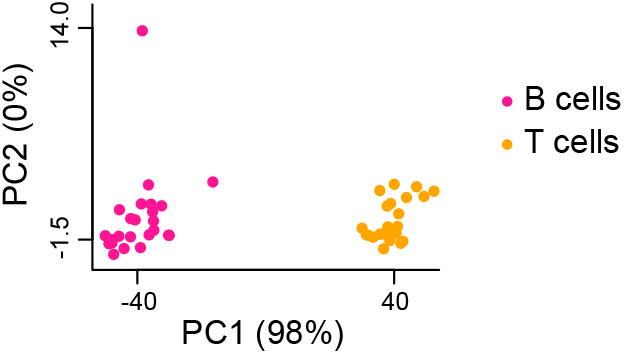
Joint principal component analysis of samples from B and T cells of KS1, KS2, and WT mice. Each point corresponds to a single sample, and points are colored according to whether they come from B or T cells. All 3 mouse genotypes (KS1, KS2, wild-type) are represented among these samples. The x and y axes correspond to the first and second, respectively, principal component; the percentage of variance each explains is indicated in the parentheses. Note that since almost all of the variance is explained by PC1, between-sample distances along the y axis are much smaller than distances along the x axis, even when they visually appear similar.

**Supplementary Figure S8.**
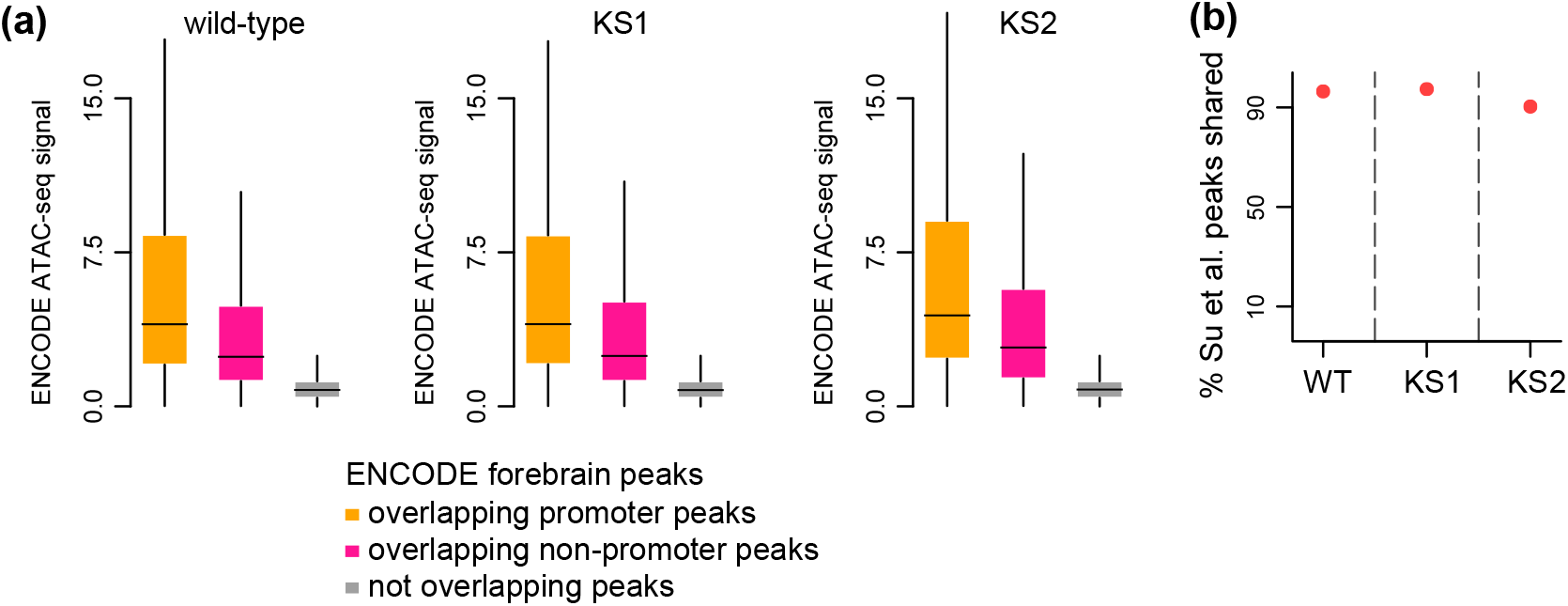
Orthogonal validation of ATAC peaks detected in KS1 and KS2 in hippocampal neurons. **(a)** The distribution of the ATAC signal (enrichment over random regions) of ENCODE peaks detected in post-natal forebrain (Gorkin et al. [53]; see also Methods), stratified according to whether they overlap peaks detected in our study within promoters, overlap peaks detected in our study outside promoters, or do not overlap peaks detected in our study. Shown for wild-type, KS1, and KS2. **(b)** The y axis depicts the percentage of ATAC peaks detected in Su et al. [66] that are also detected in neurons in our study (see also Methods), separately for each of the 3 genotypes.

**Supplementary Figure S9.**
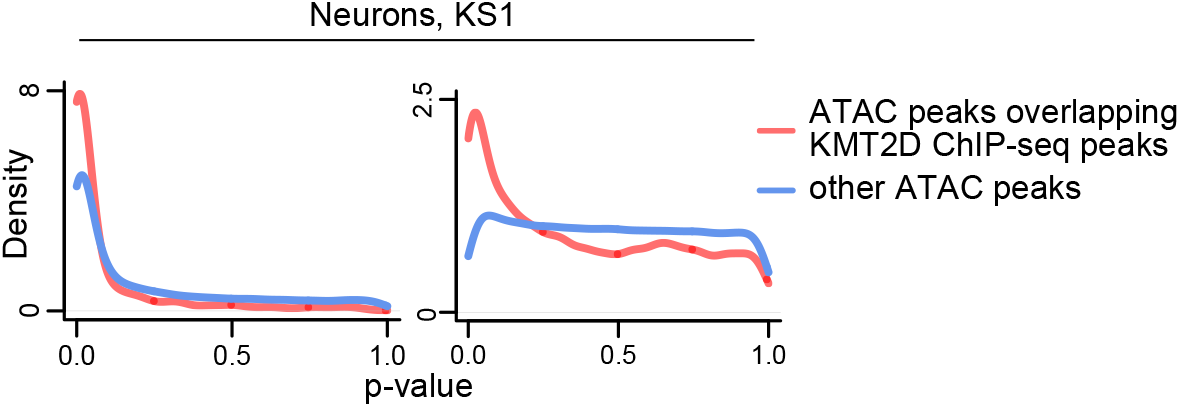
Evaluating the overlap between regions that exhibit changes in neuronal chromatin accessibility in Kabuki syndrome types 1 and regions bound by KMT2D in mouse ht22 cells. The distributions of p-values from the KS1 mutant vs wild-type differential accessibility analysis, for neuronal ATAC peaks that do and do not overlap ChIP-seq KMT2D peaks in mouse ht22 cells (a mouse hippocampal neuronal cell line). ATAC peaks in promoters and ATAC peaks outside promoters are depicted separately.

**Supplementary Figure S10.**
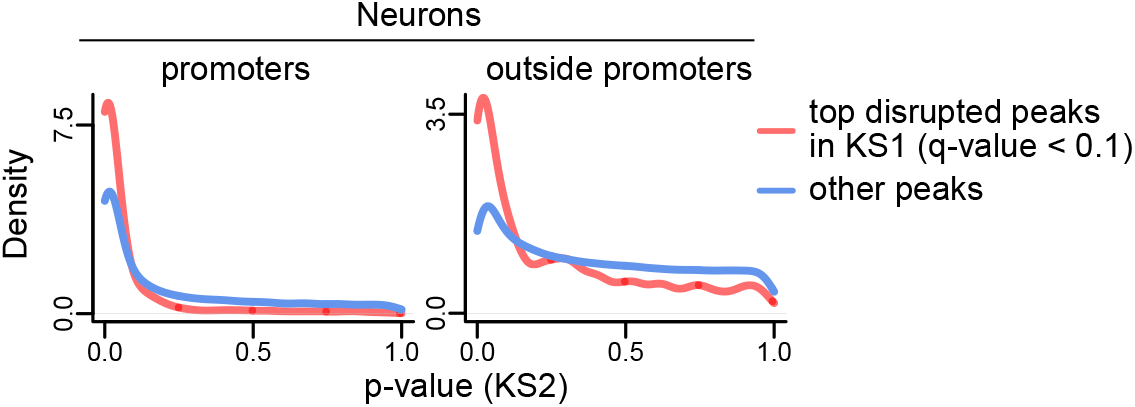
Evaluating the overlap between the regions with differential accessibility in KS1 and KS2 neurons. The distributions of p-values from the KS2 mutant vs wild-type differential accessibility analysis, stratified according to whether the same neuronal peaks are disrupted in KS1 neurons (red densities; q-value ¡ 0.1 in KS1) or not (blue densities; rest of the p-values). Shown separately for peaks inside promoters and peaks outside promoters.

### 2 List of Supplementary Tables

1. **Supplemental Table 1**: The coordinates (in mm10), log-fold changes, p-values, and q-values of the most significantly disrupted ATAC peaks (qvalue *≤* 0.1) from the KS1 vs wild-type differential accessibility analysis in neurons.

2. **Supplemental Table 2**: Like Supplemental Table 1, but for KS2.

3. **Supplemental Table 3**: The coordinates (in mm10), log-fold changes, p-values, and q-values of ATAC peaks overlapping promoters of epigenetic machinery genes from the KS1 vs wild-type differential accessibility analysis in neurons.

4. **Supplemental Table 4**: Like Supplemental Table 3, but for KS2.

5. **Supplemental Table 5**: The top 20 significant Reactome pathways based on the disruption of promoter ATAC peaks in the KS1 vs wild-type differential accessibility analysis in neurons.

6. **Supplemental Table 6**: Like Supplemental Table 5, but for KS2.

7. **Supplemental Table 7**: The gene names and ENSEMBL id’s of genes downstream of promoter ATAC peaks that exhibit disrupted accessibility in neurons, B and T cells at the 10% FDR level in KS1. If more than 1 gene are downstream of the same promoter (e.g. in cases of bidirectional promoters), all these genes are separately provided.

8. **Supplemental Table 8**: The coordinates (in mm10), log-fold changes, p-values, and q-values from the KS1 vs wild-type differential accessibility analysis in B cells of ATAC peaks that exhibit disrupted accessibility in all three cell types.

9. **Supplemental Table 9**: Like Supplemental Table 8, but from the KS1 vs wild-type differential accessibility analysis in T cells.

10. **Supplemental Table 10**: Like Supplemental Table 8, but from the KS1 vs wild-type differential accessibility analysis in neurons.

11. **Supplemental Table 11**: Like Supplemental Table 7, but for KS2.

12. **Supplemental Table 12**: Like Supplemental Table 8, but from the KS2 vs wild-type differential accessibility analysis.

13. **Supplemental Table 13**: Like Supplemental Table 9, but from the KS2 vs wild-type differential accessibility analysis.

14. **Supplemental Table 14**: Like Supplemental Table 10, but from the KS2 vs wild-type differential accessibility analysis.

